# Subtraction and division of visual cortical population responses by the serotonergic system

**DOI:** 10.1101/444943

**Authors:** Zohre Azimi, Katharina Spoida, Ruxandra Barzan, Patric Wollenweber, Melanie D. Mark, Stefan Herlitze, Dirk Jancke

## Abstract

Normalization is a fundamental operation throughout neuronal systems to adjust dynamic range. In the visual cortex various cell circuits have been identified that provide the substrate for such a canonical function, but how these circuits are orchestrated remains unclear. Here we suggest the serotonergic (5-HT) system as another player involved in normalization. 5-HT receptors of different classes are co-distributed across different cortical cell types, but their individual contribution to cortical population responses is unknown. We combined wide-field calcium imaging of primary visual cortex (V1) with optogenetic stimulation of 5-HT neurons in mice dorsal raphe nucleus (DRN) — the major hub for widespread release of serotonin across cortex — in combination with selective 5-HT receptor blockers. While inhibitory (5-HT1A) receptors accounted for subtractive suppression of spontaneous activity, depolarizing (5-HT2A) receptors promoted divisive suppression of response gain. Added linearly, these components led to normalization of population responses over a range of visual contrasts.

## Introduction

Normalization is a processing step that supports stability of transfer functions across neuronal populations despite wide ranges of external and internal input. In the visual system the necessary mathematical operations like subtraction and division were found to be supported by distinct polysynaptic local (Atallah et al., 2012; Wilson et al., 2012; Lee et al., 2012), horizontal distal (Adesnik and Scanziani, 2010a; Sato et al., 2014, 2016), and top-down cortical circuits (Angelucci and Bressloff, 2006). However, how these various intra-cortical origins are orchestrated across neuronal populations and to which extent these neuronal mechanisms can be generalized across different modalities and species remains unclear (Carandini and Heeger, 2011; Seybold et al., 2015).

Here we propose serotonergic neurons in the dorsal raphe nucleus (DRN) and their cortical modulatory effects as another origin of subtractive and divisive normalization of population activity in the primary visual cortex (V1).

Pioneering studies using electrical stimulation of the DRN revealed a general modulatory influence on evoked visual cortical responses (Gasanov et al. 1989; Moyanova and Dimov 1986). In addition, direct cortical application of 5-HT via microiontophoresis and single cell cortical recordings *in vivo* showed either suppressive or facilitative effects (Krnjevic and Phillis, 1963; Reader, 1978; Waterhouse et al., 1990). More recent studies applied antagonist and agonist of 5-HT via microiontophoresis in the monkey visual cortex, suggesting that 5-hydroxytryptamin (5-HT)-receptors are involved in normalization of visual responses (Watakabe et al., 2009; Seillier et al., 2017). However, so far, only studies on somatosensory-driven behavioral responses (Dugué et al., 2014) and in olfactory cortex (Lottem et al., 2016) specifically activated 5-HT neurons in the DRN to trigger the natural and intrinsic spatiotemporal synaptic 5-HT innervation pattern within the target cortical areas. Therefore, to investigate how DRN projections influence sensory processing directly (Davis et al., 1980), in particular, during early cortical steps of fast sensory cortical encoding (Waterhouse et al., 1990; Hurley et al., 2004; Petzold et al., 2009), optogenetic approaches were used (Ranade and Mainen, 2009; Dugué et al., 2014; Lottem et al., 2016). For example, in the somatosensory system optogenetic stimulation of 5-HT neurons in the DRN increased perceptual thresholds for tactile stimuli (Dugué et al., 2014). Interestingly, in olfactory cortex, similar optogenetic DRN stimulation reduced spontaneous (ongoing) firing without affecting amplitudes of stimulus-driven cortical responses (Lottem et al., 2016). Together, these findings suggest that the 5-HT system impacts generally on the relative weight of sensory cortical signals but may intervene in normalization of external input in a modality-specific way.

Here we used optogenetic DRN stimulation to achieve specific activation of 5-HT neurons in the DRN and to account for the presynaptic patterns of cortical 5-HT release in anesthetized mice. In particular we probed its effects on cortical population processing in the mouse primary visual cortex (V1). Further we used specific antagonists of abundant 5-HT1A and 5-HT2A receptors (Dyck and Cynader 1993; Jakab and Goldman-Rakic 1998; Leysen 2004; Shukla et al., 2014; Riga et al., 2016) in the visual cortex to disentangle contributions of these 5-HT receptor pathways to overall population activity in V1. Finally, we show how the serotonergic system mediates population response normalization, a neural operation that is supposed to be canonically implemented across sensory modalities and brain regions (Carandini and Heeger, 2011).

## Results

To control activation of 5-HT neurons in the DRN with precise timing, we used the transgenic ePet-Cre mouse line, which allows for specific expression of Channelrhodopsin2 (ChR2) in 5-HT neurons by Cre-dependent expression (Scott et al., 2005) of double-floxed adeno-associated virus (AAV) and thus, enabling real-time activation of 5-HT neurons via photostimulation (Li et al., 2005) (Figure 1). In order to simultaneously record activity of a large pool of neurons across V1, we employed wide-field optical imaging of Ca^2+^ signals which have been shown to reflect suprathreshold population activity across upper cortical layers (Wallace et al., 2008; Tian et al., 2009; Ma et al., 2014; Kim et al., 2016; Xiao et al., 2017). Specifically, we used the red-shifted fluorescent probe RCaMP (Akerboom et al., 2013; Dana et al., 2016) (Figure 1a) to minimize interference with the blue light used to activate serotonergic neurons in the DRN (Figure 1c) and to reduce light scattering in comparison to GCaMP.

**Figure 1.**
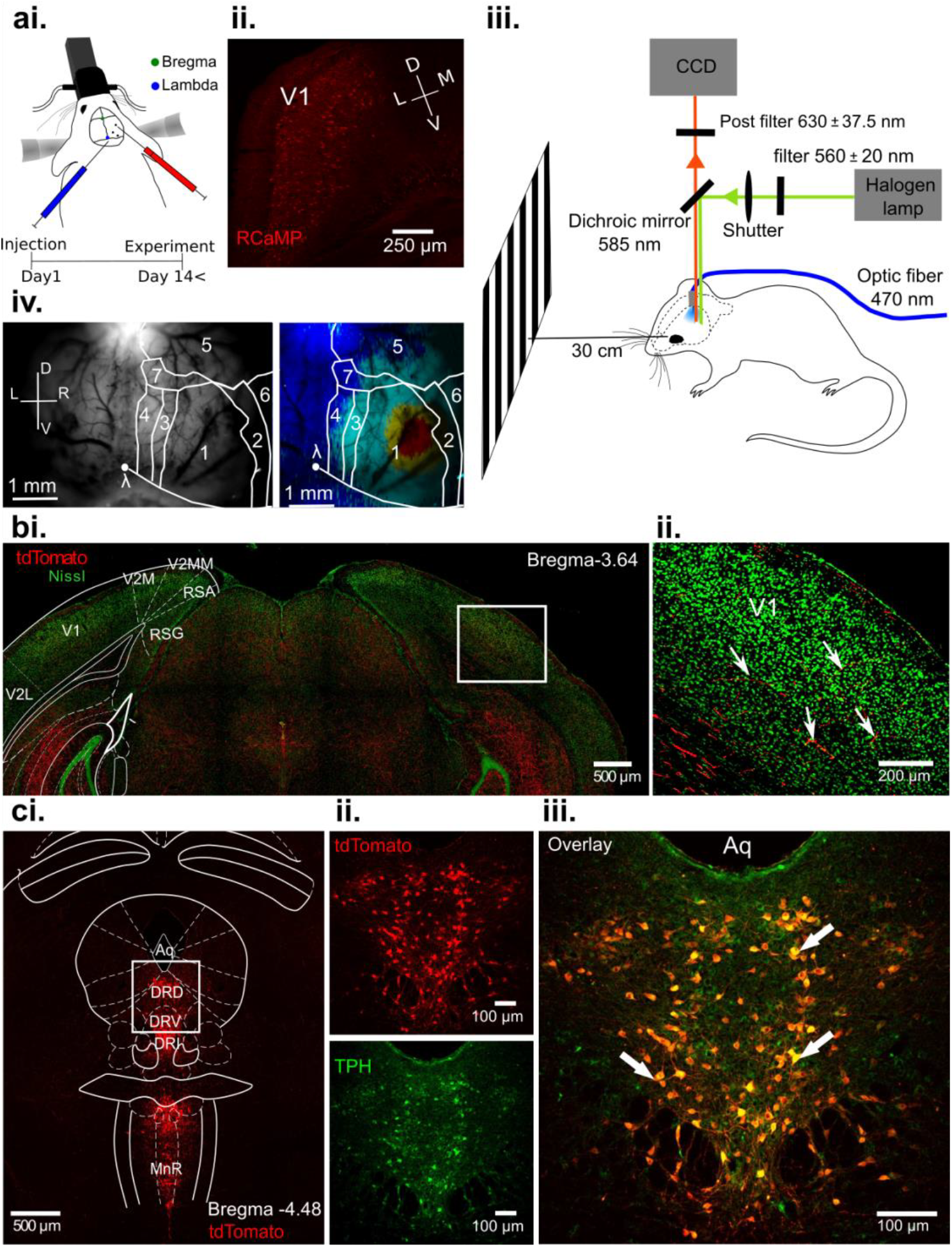
Activating 5-HT neurons in the DRN and concurrent imaging of evoked Ca^2+^ signals across primary visual cortex (V1) *in vivo*. **ai.**, Viral injections of ChR2-mCherry in DRN of ePet-Cre mice and RCaMP in V1 and S1. **aii.**, Coronal section across V1. After at least 14 days RCaMP indicator showed homogenous expression. **aiii.**, Schematic of experimental setup. DRN photostimulation was performed via an implanted optic fiber and wide-field imaging was achieved through the thinned skull (see Methods). Stimuli were displayed on a monitor at 30 cm distance to the eye contralateral to the recording hemisphere. Animals were anaesthetized and head-fixed. **aiv.**, Vascular pattern (left) of the imaged cortical region showing activation across V1 and neighboring visual areas after visual stimulation (right). 1: V1; 2: V2_L_; 3: V2_ML_; 4: V2_MM_; 5: S1; 6: A1; 7: PtA (Parietal association). **bi.**, Confocal image showing serotonergic fibers (red) co-labeled with fluorescent Nissl (green). **bii.**, Magnified image of the region outlined with white rectangle in **bi.**. Arrow heads indicate serotonergic innervation of the visual cortex. **ci.**, tdTomato fluorescence (red) at the DRN injection site. Most subnuclei which project to cortical areas including DRD, DRV, DRI and MnR show strong labeling of serotonergic cells. **cii./iii.**, Magnified view of area outlined in **ci.**. Labeling with fluorescent reporter tdTomato (**cii.**, top), antibody labeling against tryptophan hydroxylase (TPH, **cii.**, bottom), and their co-localization (**ciii.**); Aq: aquaduct.

### 5-HT influences V1 population activity through separable components of suppression

We either recorded population activity in response to visual stimuli presented repeatedly (10 times, 200 ms duration) or recorded spontaneous activity both over 30 seconds. Additionally, activity during these conditions was captured during photostimulation (train of blue light pulses at 20 Hz, 16 seconds, marked as blue bars in Figure 2aii. and 2aiv.), which was used to activate 5-HT neurons in the DRN. For each of these conditions, Figure 2a shows time courses of RCaMP signals derived from 8 different animals as mean spatial averages across V1 (30 to 50 repetitions, i.e. trials per condition). Under control conditions, i.e., without DRN photostimulation, we found activation over V1 reporting each stimulus occurrence by a rapid ramp-up of the RCaMP signal followed by slower decay towards baseline levels (Figure 2ai.). In contrast, following photostimulation (Figure 2aii.) we observed a strong suppression of evoked visual responses succeeded by later increase of the Ca^2+^ signal. To exemplify the spatial extent of these effects Figure 2b depicts image frames (500 ms time-binned) of one experiment over the first 13 seconds of recordings (black rectangles in Figure 2a, animal #8).

**Figure 2.**
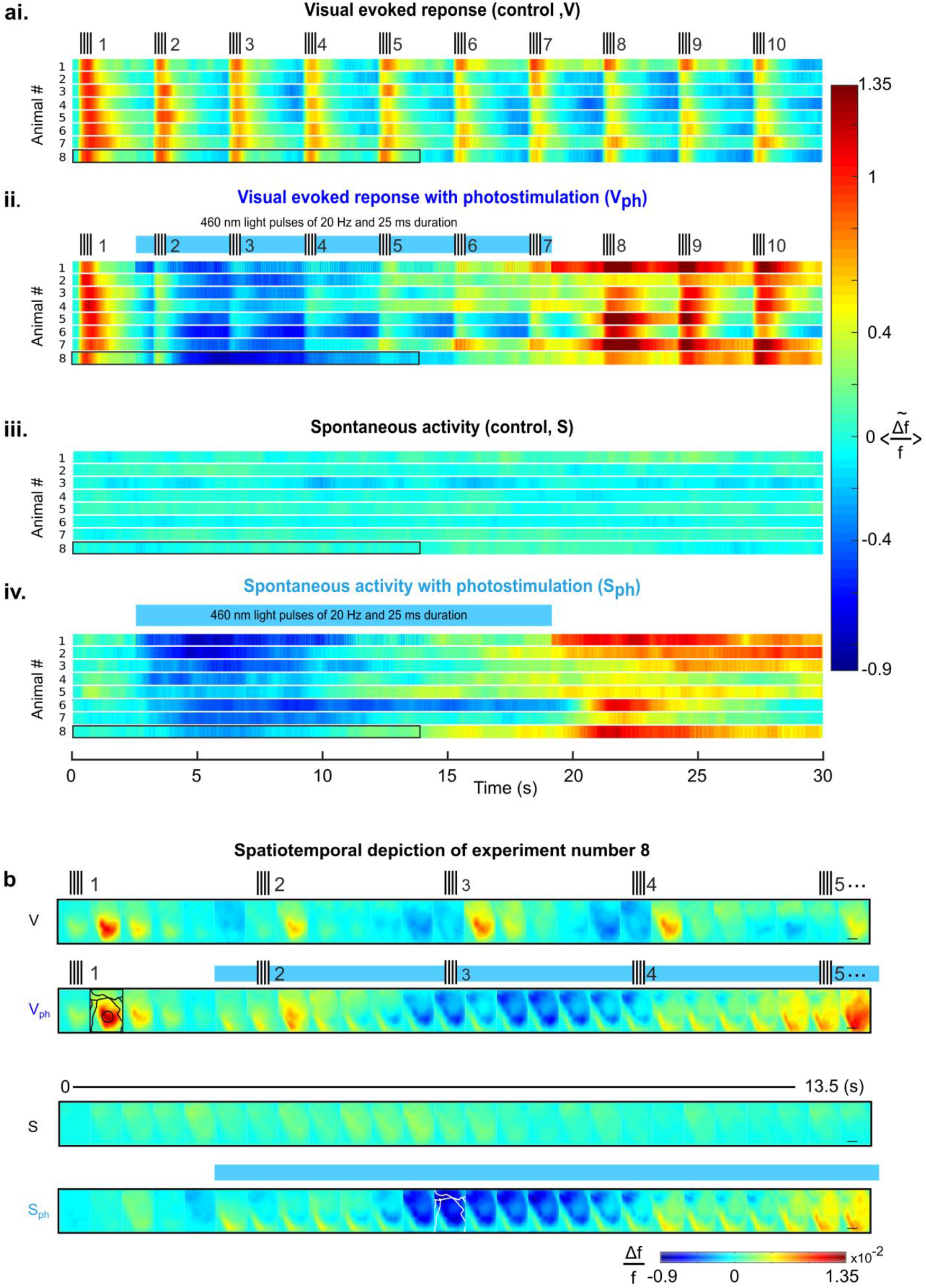
Suppression of cortical population activity after photostimulation of 5-HT neurons in the DRN. **ai.--av.**, Traces for 8 different animals as spatial averages across V1 in response to 4 different conditions. Changes in activity over time are expressed as relative change in fluorescence (Δf/f, cf. colorbars). **ai.**, Episodical visual stimulation with drifting gratings of 100% contrast (see icons on top for on/offsets). **aii.**, Visual stimulation and concurrent photostimulation of the DRN (blue bar on top marks photostimulation time). **aiii.**, Spontaneous activity. **aiv.**, Spontaneous activity and concurrent photostimulation of the DRN. **b.**, Example experiment (#8, encircled black in above traces) depicting 13 seconds of recording across the cortical surface before spatial averaging (horizontal black lines in right frames delineate 1 mm). Rows of image frames (500 ms time binned) correspond to conditions as shown in **a.**. Note that 5-HT-induced suppression encompasses the entire imaged cortical region. Different cortical areas (as specified in Figure 1a.) are outlined black and white (see frames within row two and four, respectively). Small signal increases at lower left corners of image frames indicate artifacts due to partial interference with photostimulation light.

Importantly, 5-HT-mediated suppression of cortical activity was also present without external visual input. That is, following photostimulation, ongoing spontaneous cortical activity rapidly declined below baseline levels (Figure 2aiii. and 2aiv.). Such suppression included cortical areas beyond V1 (white contours in Figure 2b, 4^th^ row; for assignment of cortical areas see Figure 1aiv.), suggesting 5-HT effects on spontaneous drive through widespread ascending projections from the DRN across the entire cortex (Hale and Lowry, 2011).

Figure 3 summarizes these results as time traces (mean across animals, n = 8) over the whole period of recordings. Responses to visual stimuli under control conditions were characterized by peaks of activity with small adaptive decrease in amplitudes and baseline over time (Figure 3ai., black trace). Upon photostimulation of 5-HT neurons in the DRN (blue area marks photostimulation duration), visually-evoked responses markedly decreased in amplitude (Figure 3aii., dark blue trace; for sham controls see Figure 3—figure supplement 3-1). This reduction in the gain of the responses appeared independent from the decline in baseline (i.e., activity levels before each stimulus onset). Instead, the baseline envelope of these traces displayed a similar time course as observed for 5-HT-induced suppression in spontaneous activity devoid of visual input (Figure 3aiii., light blue trace). To further assess how much of the suppression in evoked amplitudes was due to suppression observed during spontaneous drive, we subtracted image frames obtained under spontaneous conditions from those under evoked conditions both during DRN photostimulation. This computation was done pixelwise and across single trials in each experiment. The suppression of evoked activity remained clearly visible in the resulting trace (Figure 3aiv.) suggesting that the suppression in response amplitudes was independent of the 5-HT-induced reduction of the spontaneous drive. To validate these findings, we sampled response amplitudes by averaging over the time span during which activity was >50% of its maximum amplitude (time window w_2_ in Figure 3bi. inset) and subtracted average baseline levels 200 ms before each stimulus onset (time window w_1_ in Figure 3bi. inset). This calculation provides discrete, baseline-independent, amplitude values for each single stimulus event (“Stim #1-10”) across evoked conditions, plotted in Figure 3bi as mean ± 2 s.e.m. over all experiments. Photostimulation-induced suppression of evoked amplitudes was highly significant as compared to the control condition (Figure 3bi., cf. dark blue and black curve, respectively) and approached initial values thereafter. The amount of suppression remained similar after subtraction of the spontaneous drive during photostimulation (Figure 3bi., gray trace), verifying that modulation of response-gain was restricted to the evoked component. In contrast, the time course of the baseline component in the evoked responses during photostimulation was analog to the time course obtained for photostimulation during spontaneous drive (Figure 3bii., cf. dark blue and light blue traces), characterized by initial suppression followed by the above-mentioned later rise of the RCaMP signal (the nature of this later rise is further elucidated in Figures 4 and 5). Altogether, we identified two suppressive components independently shaping cortical activity through photostimulation of 5-HT neurons in the DRN, one driving baseline levels and one affecting the amplitude (i.e., the gain) of visually evoked responses.

**Figure 3.**
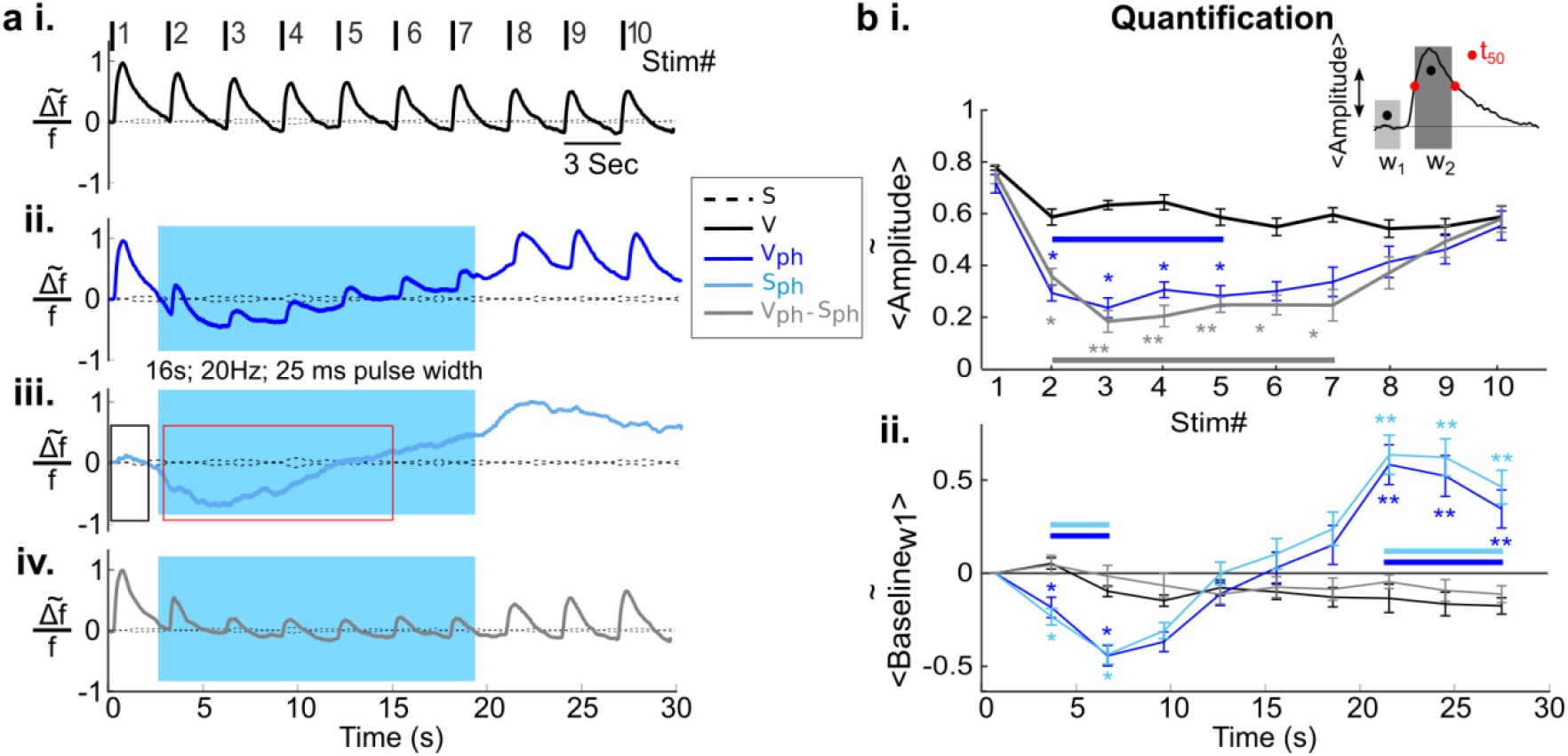
Serotonergic input affects different population response components. **ai.**, Control condition: Spatial averages across the center of activity in V1 (see black circle in Figure 2b, 2^nd^ frame 2^nd^ row, for an example) after visual stimulation (numbered above traces). **aii.**, Same as **ai.** with concurrent DRN photostimulation (16 s; 20 Hz; 25 ms pulse width; on/offset marked as blue rectangle in the background). **aiii.** Spontaneous component: Same as **aii.** during spontaneous activity, i.e., without visual stimulation. Black and red rectangles sketch time windows used for quantification of spontaneous component suppression (cf. Figure 5**cii**.). **aiv.**, Evoked component after subtraction of the spontaneous component shown in **aiii.**. **bi.**, Quantification of evoked response amplitudes. Amplitudes were calculated by subtracting average baseline levels 200 ms before each onset of visual stimulation (w_1_) from the mean level during which peak activity was at half maximum (w_2_, see inset). Black: Controls; dark blue: Visual responses during concurrent DRN photostimulation; gray: Visual stimulation and concurrent DRN photostimulation after subtraction of spontaneous component. Bars matching color of the traces mark time segment with *P* = 0.016 (blue) and *P* = 0.008 (gray) after run-length correction test (see Methods). **bii.**, Time course of baseline activity (w_1_). Colors denote same conditions as shown in legend **a.**. Bars mark additional run-length tested segments for both dark and light blue traces, *P* = 0.012 (dark blue) and *P* = 0.011 (light blue). Note the offset between the blue curves, reflecting a continuous small adaptive decrease in baseline observed under visually evoked conditions (black and gray traces) regardless of the presence of 5-HT (cf. 3**ai.** and **3aiv.**). All graphs depict mean values across 8 animals, error bars indicate 2 s.e.m.. \*\**P*<0.01, \**P*<0.05, two-sided t-test.

**Figure 4.**
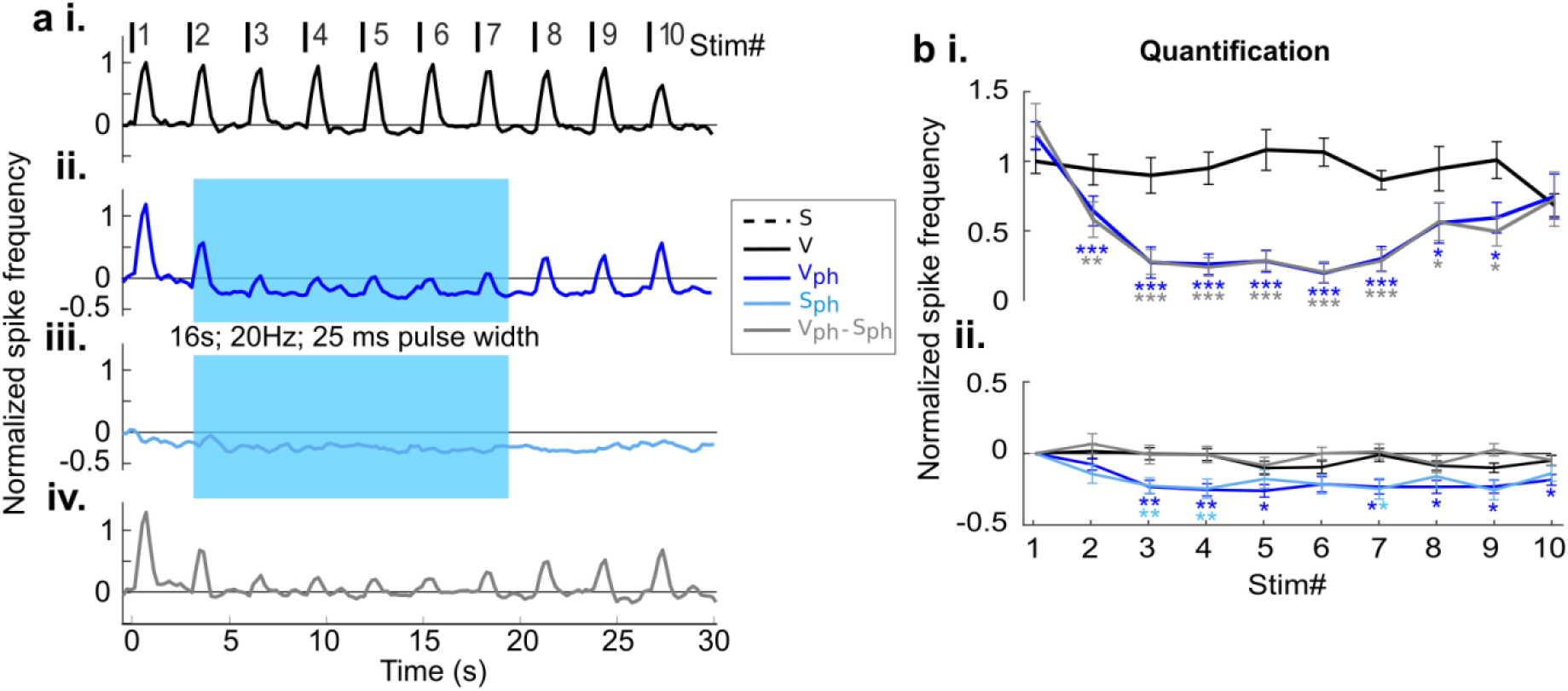
Extracellular recordings across different layers confirm observed suppressive effects in V1. **ai.**, Control condition: Visually evoked responses without DRN photostimulation. Data were normalized to peak activity in response to the first visual stimulus. Time periods of visual stimulation marked with black bars on top. Mean across 6 mice (20 different recordings at cortical depths between 250 and 750 μm, each comprising 10-20 trials) **aii.**, Same as **ai.** with DRN photostimulation (blue background area depicts time of photostimulation, same as in Figure **3aii.**). **aiii.**, DRN photostimulation during spontaneous activity. **aiv.,** Evoked component after subtraction of the spontaneous component shown in **aiii.**. **bi.**, Quantification of evoked response amplitudes. Same calculations as used for Figure **3bi.**. Black: Controls; dark blue: Visual responses during concurrent DRN photostimulation; gray: Visual stimulation and concurrent DRN photostimulation after subtraction of spontaneous component. **bii.**, Time course of baseline activity. Colors denote stimulation conditions as shown in **bi.**. Light blue trace: Time course of spontaneous activity during DRN photostimulation. All graphs depict mean values, error bars indicate 2 s.e.m.. \*\*\**P*<0.001, \*\**P*<0.01, \**P*<0.05, two-sided t-test.

**Figure 5.**
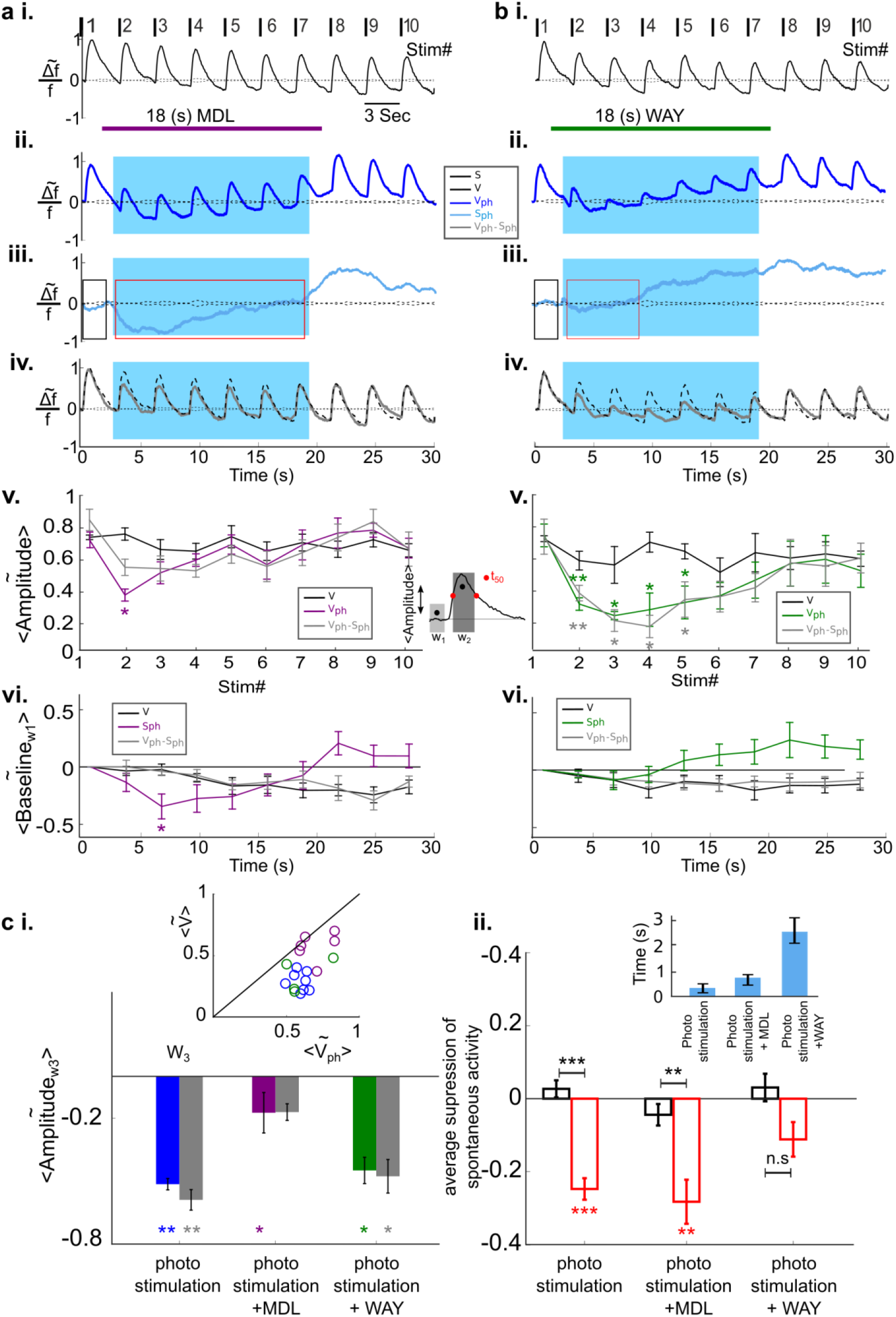
5-HT2A and 5-HT1A receptors act synergistically on different components of suppression in V1. **ai.-iv.**, Same conditions as in Figure 3**ai.-iv.** with additional blocking of 5-HT2A receptors via microiontophoresis of MDL (purple horizontal bar shows timing of MDL administration, 18 s, average across 6 animals). Dashed trace in **aiv.** repeats control trace shown in **ai.**. **av.** and **avi.** depict quantification of traces shown in **ai.-iv.** for amplitude and spontaneous components, respectively. Amplitude values were calculated in the same way (see inset in **av.** for amplitude calculation) as for the experiments without drug application (cf. Figure 3 **bi-ii.**). **bi.-vi.**, Same as **ai.-avi.** using WAY for blocking of 5-HT1A receptors (green horizontal bar shows the timing of WAY administration, 18 s, average across 4 animals). **ci.**, Summary of amplitude differences to control condition (V). Quantified amplitude values were averaged over visual responses during the time span photostimulation was applied (Stim #2 to #7); color scheme as above; inset depicts single experiments. **cii.**, Summary of changes in spontaneous activity levels. In each experiment values were averaged across time windows during photostimulation in which activity levels were significantly below baseline (onset of window) until reaching values above baseline (offset of window), as sketched by red rectangles in Figures 3**aiii.**., 5**aiii.**, and 5**biii.**. These mean values were subtracted from average activity in the time window before photostimulation onset (black rectangles); bar colors match time windows. Inset: Point in time after onset of DRN photostimulation at which spontaneous activity changed significantly from baseline (320 ± 5 ms without blockers, left bar). All graphs depict mean values, error bars indicate 2 s.e.m.. \*\*\**P*<0.001, \*\**P*<0.01, \**P*<0.05, two-sided t-test.

To investigate whether the observed suppressive effects reflect decrease in spiking output of V1, we recorded extracellular multi-unit activity (MUA) of pyramidal cells using the same experimental paradigms as employed in our wide-field RCaMP recordings (Figure 4a; 20 recordings of MUA in 6 animals). Similar to the imaged RCaMP signals we found a suppressive component affecting the gain of evoked responses (Figure 4aii. and aiv.) as well as a component reflecting suppression of baseline activity (Figure 4aiii.; for an example showing single trials as raster plots see Figure 4—figure supplement 4-1). These suppressive effects on spiking activity were found similarly across different cortical depth (z = −0.8729, *P* = 0.3827, Wilcoxon rank sum test). Note that in contrast to the imaged RCaMP signal, spike recordings showed suppression throughout the entire time span of photostimulation without a later rise of amplitudes above baseline values (cf. times after offset of photostimulation). Thus, the observed late increase in RCaMP signals most likely reflects increase of intracellular Ca^2+^ levels associated with 5-HT receptor-mediated activation of the Gq/11 pathway (Jang et al., 2012; Millan et al., 2008) and concomitant activation of store-operated channels (Celada et al. 2013). Figure 4b shows a quantification of the spike recordings using similar analysis as applied for the imaging data shown in Figure 3b. Baseline-independent suppression of the gain of evoked spike responses was highly significant (Figure 4bi.) as well as the initial suppression of the spontaneous component (Figure 4bii.). Altogether, the high similarity obtained for spike recordings and RCaMP imaging indicates that the observed suppressive signals report transient suppression of suprathreshold population activity in V1.

### Isolation of 5-HT2A and 5-HT1A receptor contribution to suppression

We next investigated whether the observed different suppressive components were mediated via activation of different 5-HT receptor pathways (Leysen, 2004; Santana et al., 2004; Hannon and Hoyer, 2008). After specifically blocking 5-HT2A receptors via microiontophoresis of MDL (see Methods) and parallel photostimulation of the DRN, we found response amplitudes only slightly reduced (Figure 5aii.). Thus, after subtraction of the baseline component, evoked amplitudes during activation of 5-HT neurons in DRN were nearly identical to those under control conditions (cf. gray and stippled black traces in Figure 5aiv.) as soon as the 5-HT2A antagonist becomes effective (Figure 5av., compare purple with black and gray traces; note that the delayed onset of the drug effect is likely due to limitations in the sphere of influence of microiontophoretically applied substances). In contrast, the suppression of the baseline component remained present (Figure 5aiii., light blue trace) and was similar to DRN photostimulation during spontaneous activity without drug application (Figure 3aiii.). In addition, the RCaMP signal remained negative during photostimulation (Figure 5avi., purple trace) as compared to conditions without drug application (Figure 3bii. light blue trace), suggesting reduction of the aforementioned intracellular Ca^2+^ accumulation by effectively blocking 5-HT2A receptors. Conversely, blocking of 5-HT1A receptors by microiontophoresis of WAY (see Methods) abolished suppression of baseline activity in both spontaneous and evoked conditions (Figures 5biii. and 5bvi., light blue and green trace, respectively), while suppression of response amplitudes was not affected (Figures 5bii., biv., and 5v.). Altogether, these results establish that 5-HT1A and 5-HT2A receptors target two different components of cortical activity. Whereas activation of 5-HT2A receptors dominantly serves suppression of evoked amplitudes (Figure 5ci.), activation of 5-HT1A receptors initiates suppression of spontaneous baseline activity (Figure 5cii.). Together, both components influence visual processing within a few hundred milliseconds after activation of 5-HT neurons in the DRN via photostimulation (Figure 5 cii., left bar in inset).

### 5-HT-induced normalization of V1 population responses to visual contrast

To explore how 5-HT-induced suppression affects cortical response normalization, we presented visual stimuli at varying contrast. Responses were imaged under control conditions interleaved with trials in which the DRN was photostimulated 500 ms before the onset of the visual stimulus, to ensure that 5-HT input affected the evoked responses (cf. Figure 5 cii., inset and legend). Control responses progressively declined in amplitude with decreasing contrast, as expected (Porciatti et al., 1999) (Figure 6ai). However, upon photostimulation of the DRN, evoked responses were markedly decreased for all contrasts (Figure 6aii.). We accounted for possibly underlying normalization effects on response amplitude under serotonergic influence in two steps. First, we subtracted the baseline component observed for the spontaneous condition during photostimulation (light blue traces, S_ph_ in Figure 6aii and aiii.) from each of the different contrast conditions, unraveling modulation in the gain of the evoked visual responses independent of the baseline suppression (Figure 6aiii). Next, we further normalized responses by divisive scaling to the maximum amplitude at 100% contrast (Figure 6aiv.). After these two simple operations of normalization, we obtained a close match between traces of controls and of traces during activation of 5-HT neurons in the DRN (compare Figure 6ai. and 6aiv.). Figure 6b depicts amplitude values (for quantification of values see Figure 3bi. inset) of all traces shown in Figure 6ai.-aiii. The parallel slope of the curves for controls (black) and photostimulated conditions after subtraction of the spontaneous component (gray) indicates equal gain control across different contrasts in both conditions. Note that the serotonergic modulation of the gain of amplitudes became increasingly independent of the baseline component with increasing contrast, as indicated by the diverging blue and gray curves (Figure 6b.).

**Figure 6.**
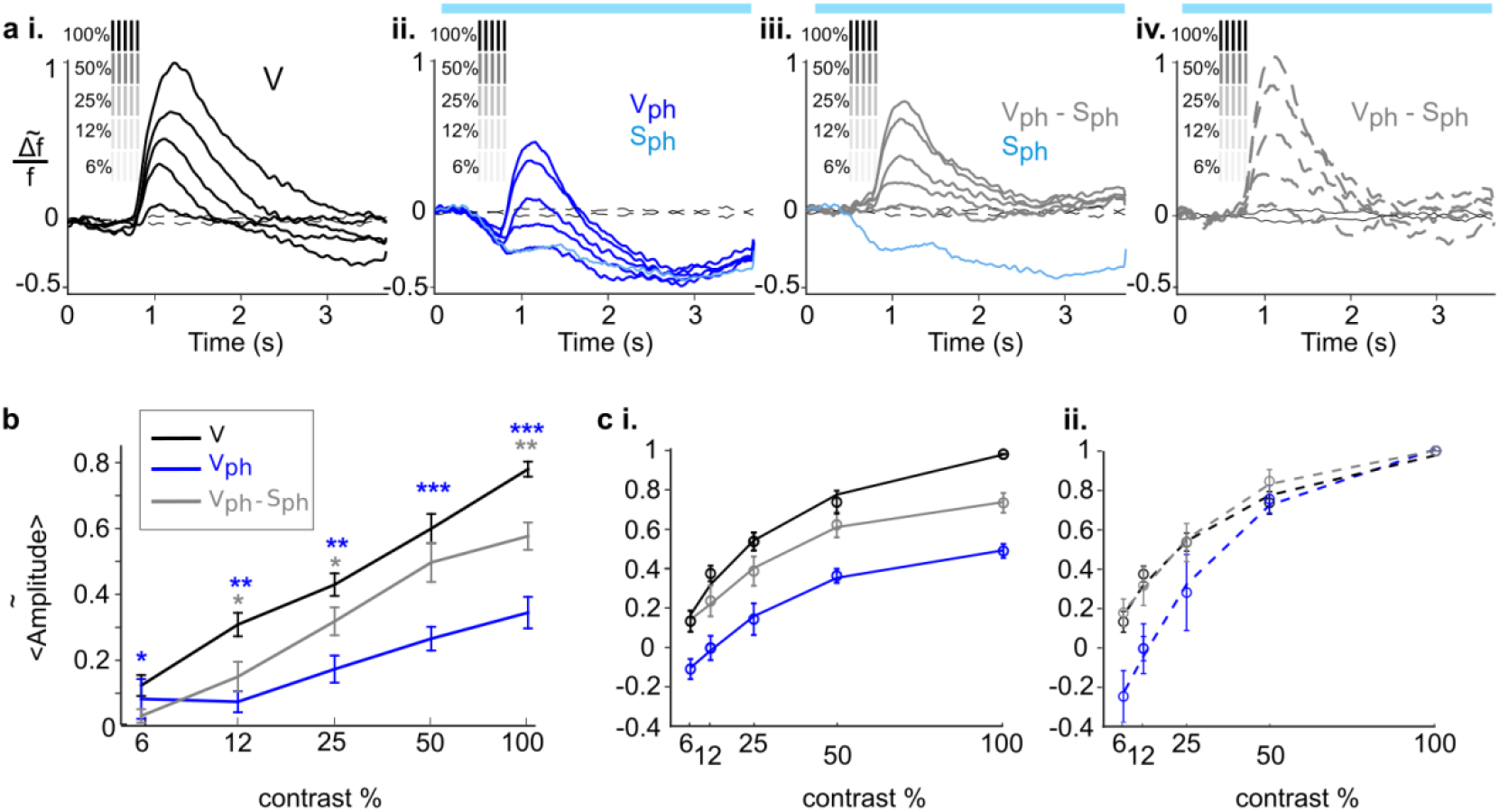
Subtractive and divisive components provide normalization of visual cortical responses during photostimulation of 5-HT neurons in the DRN. **ai.**, Control conditions (V): Time courses of activity (spatial averages across V1) after visual stimulation with gratings of different contrast (n = 7 mice). Values were scaled to maximum amplitude at 100% contrast. Grating icons identify timing of visual stimulation and corresponding contrast. Black dashed lines show baseline activity without visual stimulation. **aii.**, Responses to visual stimulation at variable contrasts and concurrent DRN photostimulation (V_ph_); values were scaled to control traces shown in **ai**. Light blue trace: Spontaneous component, i.e., response to DRN photostimulation during spontaneous activity (S_ph_). **aiii.**, Evoked component, i.e., responses after removal of the spontaneous component (light blue trace) during DRN photostimulation (i.e., V_ph_-S_ph_). **aiv.**, Divisive normalization: Evoked component (gray traces in **aii.**) was scaled to maximum amplitude at 100% contrast. **b.,** Amplitude values as a function of contrast (see main text and legend of Figure 3 for calculation); same color code as in **a**. **ci.**, Amplitude values shown in **ai.-aiii.** fitted to the Naka-Rushton function (see Method) (R^2^: 0.84, 0.95 and 0.91 for black, blue and gray, respectively). **cii.**, Values shown in **ci.** after two steps of normalization. For reasons of better comparison, black curve in **ci.** is replotted as stippled line together with single values. Blue curve depicts divisive normalization without prior subtraction of the spontaneous component. Gray curve depicts normalized peak maxima after subtraction of the spontaneous component. Note the close overlap between black stippled curve and gray stippled curve, indicating two-component (subtractive and divisive) normalization of contrast responses (R^2^: 0.82, 0.94 and 0.92 for black, blue and gray, respectively). Blue horizontal bars mark time of DRN photostimulation. All graphs depict mean values (n = 7 mice), error bars indicate 2 s.e.m.. \*\*\**P*<0.001, \*\**P*<0.01, \**P*<0.05, two-sided t-test.

In addition, in comparison to control conditions, we found that DRN photostimulation induced significant decrease in the duration of population responses across all contrasts (Figure 6—figure supplement 6-1iii.)

To further evaluate how serotonergic gain control affects normalization over varying contrast, we fitted amplitude values (mean across experiments) obtained for each contrast to the Naka-Rushton function (Figure 6ci.). Applying this function to the above 2-step normalized values revealed high similarity to controls (Figure 6cii., cf. gray and black stippled curves). Importantly, pure divisive normalization (i.e., without considering the subtractive component in a first step, blue stippled curve in Figure 6cii.) failed to replicate the contrast response function of controls, particularly at lower stimulus contrast.

In sum these results confirm the existence of two independent, but jointly operating components involved in 5-HT-induced visual response normalization: One component acting subtractive on baseline and the other acting divisively on response gain, each controlled by different 5-HT antagonistic receptor pathways.

## Discussion

Changes in 5-HT levels are known to modulate cortical state, affecting all sorts of higher cognitive function engaged in perception and decision (Doya, 2008) such as emotion, memory, or expectation of reward (Ranade and Mainen, 2009), and thus, its long-term malfunction facilitates psychiatric disorders like anxiety, depression and schizophrenia (Soubrié, 1986; Lucki, 1998; Geyer and Vollenweider, 2008; Zhang and Stackman, 2015; Urban et al., 2016). Here we showed that activation of 5-HT neurons in the DRN causes visual input normalization across neuronal populations in V1 through rapid suppression of two entities of cortical activity — internally generated (spontaneous baseline) and stimulus-driven (sensory evoked) activity.

How may the 5-HT-mediated computation of visual response normalization be implemented at the cortical circuit level? Local administration of pharmacological blockers revealed that the bulk of these effects was attributable to the joint activation of antagonistic (inhibitory) 5-HT1A and (excitatory) 5-HT2A receptors that most likely achieve synergy by operating predominantly on antagonistic subpopulations of neurons (i.e., pyramidal vs. GABAergic). The identified subtractive component is likely due to activation of 5-HT1A receptors leading to direct suppression of pyramidal cell output through decrease in synaptic excitation. Indeed, hyperpolarization of excitatory neurons was recently shown as an additional source of normalization processes (Sato et al., 2016). In contrast, the 5-HT-induced divisive component, acting on population response gain, is most likely conveyed by predominant activation of 5-HT2A receptors in GABAergic neurons, simply because 5-HT2A couples to the Gq/11 pathway (Hannon and Hoyer, 2008) leading to an increase in neuronal firing rather than suppression. Thus, suppression must partly be mediated disynaptically. Furthermore, this proposition fits to the existing literature in two mutually reinforcing key aspects. First, divisive modulation of visual cortical responses was shown to be specifically mediated by soma-targeting parvalbumin (PV) interneurons (Wilson et al., 2012, but see Lee et al., 2012; Seybold et al., 2015). Second, using immune-histochemical and electrophysiological approaches in genetically modified mice, it could be demonstrated that, in the case of interneurons, 5-HT2A receptors are dominantly expressed in the subclass of PV cells (Weber and Andrade, 2010; see also Figure 6—figure supplement 6-2). Importantly, such PV cell-mediated hyperpolarization does not rule out a concomitant increase in depolarizing currents through activated 5-HT2A receptors in pyramidal neurons (Figure 6-figure supplement 6-2), overall producing shunting inhibition and resulting in increased conductance. Such mechanism is known to not only affect the gain but also the time constant of neuronal responses (Carandini and Heeger, 2011). Consistent with this idea we found that response-onset latency decreased while response duration increased as a function of stimulus contrast in both control and photostimulated conditions (Figure 6-figure supplement 6-1ii.). Additionally, however, in comparison to controls, the duration of responses was significantly reduced during photostimulation of the DRN (Figure 6-figure supplement 6-1iii.). Briefer neuronal responses reduce the time span for temporal summation and, in turn, reduce response amplitudes (Mante et al., 2008). Thus, the additional reduction in response duration under influence of 5-HT may indicate involvement of shunting inhibition in pyramidal neurons to effectively controlling response gain.

There are some considerations with respect to the interpretation of our results that should be considered. First, a recent study suggested that wide-field Ca^2+^ signals correlate more strongly to neuronal activity in the neuropil than with activity of neurons in cortical layers 2/3 (Allen et al., 2017). However, our multi-unit spike recordings of pyramidal cells were highly consistent with the recorded RCaMP signals and revealed similar effects across superficial and deeper cortical layers. Thus, we are confident that the RCaMP signals which we report reflect spiking activity at the adjusted focal depth (Wallace et al., 2008; Kirmse et al., 2015; Xiao et al., 2017). Second, we used wide-field RCaMP imaging to capture net effects across large populations of neurons across V1. This is of advantage because sensory processing is mediated by neuronal populations (Dayan and Abbott, 2001). On the downside, with this technique distinct microcircuit mechanisms are not resolved, which can be quite diverse across populations of interneurons (Seybold et al., 2015) and across different cortical layers (Adesnik and Scanziani, 2010). Third, our optogenetic stimulation activates specifically 5-HT neurons in the DRN but uniformly. Thus, considering the broad topographic organization within the DRN (Muzerelle et al., 2016) unspecific influence of other cortical and subcortical areas modulated by activation of 5-HT neurons cannot be ruled out as these areas may additionally influence V1 or back project to DRN. On the other hand, our optogenetic approach provides stimulation of the serotonergic system in a closed-loop, thus, including all brain structures involved in the observed suppressive effects within V1. Finally, a previous study in monkey V1, using single cell recordings and pharmacological 5-HT1B and 5-HT2A receptor activation via microiontophoresis in anaesthetized preparations, revealed effects on both spontaneous activity and response gain (Shimegi et al., 2016). In contrast to ours and the above study, where the additional subtractive component essentially contributed to contrast-gain control at lower contrast, a more recent study in monkey V1 found, however, for the majority of recorded cells only divisive scaling of evoked responses (Seillier et al., 2017). As their recordings were done in awake monkeys, these differences may indeed reflect principal changes in cortical state and 5-HT levels during wakefulness (Portas et al., 2000; Tyree et al., 2017) and/or species-specific differences in individual 5-HT receptor sets (“receptomes”) tailored to task-specific demands on signal integration. Alternatively, because 5-HT was applied by microiontophoresis in their study, the difference may result from difficulties in determining baseline stationarity or from the inevitable lack of stimulating layer- and cell type-specific synaptic weight of DRN synapses associated with local iontophoretic 5-HT application, a limitation that we circumvented using optogenetic stimulation.

In summary our data suggest 5-HT neurons in DRN as a powerful source of the neuromodulation of visual cortical responses. On the one hand it may be not too surprising to find serotonergic receptors implicated in normalization, as this is true also for other modulatory transmitter systems (Doya, 2008; Dayan, 2012; Pinto et al., 2013). Given their adequate trafficking across neuronal compartments and types of neurons, 5-HT receptors simply fulfill the needed temporal properties (Roerig and Katz, 1997) to act within established canonical normalization circuits. However, here we newly identified divisive and subtractive signals arising from a distinct origin (DRN) that constitutes a relay of a diversity of cortical and subcortical projections (Pollak Dorocic et al., 2014). Worthy of note, the DRN receives also projections from the retina (Huang et al., 2017), raising the possibility that normalization signals arising from rapid and sustained changes in visual input can, in addition to other parallel pathways, directly be passed on to the cortex.

## Material and methods

All experimental procedures were carried out in accordance with the European Union Community Council guidelines and approved by the German Animal Care and Use Committee in accordance with the Deutsches Tierschutzgesetz and the NIH guidelines. Adult ePet-Cre (Scott et al., 2005) mice (both sexes) were used. After preparatory surgery mice were housed individually and kept in 12 hours light/dark cycle with food and water ad libitum.

### Viral injections and implant of optical fiber

Cre-dependent AAV [AAV9.EF1.dflox.hChR2(H134R)-mCherry.WPRE.hGH] was injected into the DRN of ePet-Cre transgenic mice (Figure 1ai.), and viral construct of the red-shifted calcium indicator RCaMP [AAV1.syn.jRCaMP1a.WPRE.SV40] (Akerboom et al., 2013) was injected into visual and somatosensory cortex based on stereotactic coordinates (Franklin and Paxinos, 2001). All the viruses were obtained from the University of Pennsylvania (100 μL at titer ≥ 1×10^13^ vg/mL).

Animals were anesthetized with isoflurane (4% induction and 2% for maintenance) via a nose mask and received 0.5 ml subcutaneous bolus of isotonic 0.9% NaCl solution mixed with Buprenorphine (2 μg/ml) and Atropin (3 μg/ml). To maintain body temperature a heating pad (37°C) was placed below the animal during surgery and experiment. Before sagittal incision along the midline, 2% Lidocaine was applied to provide additional local anaesthesia. The skull was thinned until surface blood vessels became clearly visible. The DRN was localized with the aid of stereotactic coordinates (Franklin and Paxinos 2001; Lowery et al. 2010). A small craniotomy was made (−4.7 Anteroposterior (AP) and 0 mediolaterla (ML) to Bregma) and a customized glass pipette attached to a 20-ml syringe was lowered into the brain to a depth of 2.5 mm below the brain surface using a micromanipulator. Viral solution containing ChR2 construct was delivered via small pressure injections (100 μm steps upwards until the depth of 1.7 mm, with an injection interval of 5 minutes). After injections a custom made optical fiber (200 μm, 0.37 NA, Thorlabs) attached to a ceramic ferrule (Thorlabs) was implanted at tissue depth of −1.5 mm dorsoventral (DV) and 0.5 mm anteroposterior (AP) to lambda. After implantation, the ferrule was secured to the skull with transparent dental cement (Super Bond C&B set, Hentschel-Dental). Viral solution containing RCaMP construct, was injected at two locations in visual cortex (−4.2 AP, 2.5 ML, and −3.5 AP, 2 ML to bregma) and at one location in somatosensory cortex (−1.5 AP, 2.5 ML to bregma). At each of these cortical injection sites 0.5 μl of viral solution was delivered at depth of ~600 μm and 300 μm in four steps with 10 minutes intervals between each injection step. The thinned and exposed skull was covered with transparent dental cement and nail polish. Finally, a head holder was attached (Pi-Ku-Plast HP 36, Breedent) to the skull in order to provide a clear and easy accessible imaging window for chronic experiments. Control animals matching genetic background and age received either injections with 0.9% NaCl solutions or none.

### Visual stimuli and DRN photostimulation

For photostimulation of serotonergic neurons in the DRN, pulses of blue light (470 nm, using a LED driver and emitter system, Plexon) was delivered via an optical fiber attached to the implant at a frequency of 20 Hz (25 ms pulse width). Such a frequency has been shown to evoke a robust increase of activity in the DRN above the baseline (Dugué et al., 2014). Photostimulation started 2.5 s after the onset of recording and lasted for 16 seconds. The power of light at the tip of the fiber was ~1 mW.

Vertical square-wave gratings (0.04 cycle/deg) moving at 2 Hz were presented on a monitor (100 Hz, mean luminance 40 cd/m^2^, Sony Triniton GDM-FW900, Japan) with its center placed 30 cm away from the eye that was contralateral to the cortical recording site, overall covering ~40 × 60 deg of the visual field. Eyes were covered with semipermeable zero power contact lenses to prevent drying-out or corneal edema. Each experiment comprised 25-40 trials (i.e., repetitions of stimulus conditions). Each trial consisted of four different conditions presented in pseudorandom order: 1) Blank condition, during which a uniform isoluminant gray screen was shown. These recordings were repeated twice within a trial and served as a measure of spontaneous activity. Blanks were also used to calculate relative changes in fluorescence (Δf/f, see Data analysis). 2) Visually evoked condition (controls), during which moving gratings were episodically (10 times) presented with duration of 200 ms at an interval of 3 s (during which the blank was presented). 3) Visually evoked condition, during which the DRN was photostimulated. 4) Spontaneous condition (blank) during which the DRN was photostimulated. Each condition lasted 30 s (including 200 ms prestimulus time) and time interval between conditions was 60 seconds.

In the set of experiments where different visual contrasts were used (100, 50, 25, 12.5, 6.25%), a grating stimulus was presented once for 200 ms (starting 700 ms after onset of recording). In conditions comprising additional photostimulation, its onset was at 500 ms and lasted until end of each trial (3.7 seconds in total). Additionally, spontaneous activity with and without photostimulation was recorded with identical timing to conditions comprising a grating stimulus. All conditions were presented in pseudorandomized order (interstimulus interval was 60 seconds).

During recordings mice were kept under mild anaesthesia (0.5-1% isoflurane) delivered via a nose mask.

### Imaging of fluorescent RCaMP signals

Image frames (1 pixel covered ~67 μm of cortical surface followed by additional 3 × 3 binning online) were collected at a rate of 100 Hz using an Imager 3001 system (Optical Imaging Inc, Mountainside, NY). The camera was focused ~300 μm below cortical surface. To record changes in fluorescence of the RCaMP indicator, the brain was illuminated with excitation wavelength of 520-560 nm. Emission light >585 nm was collected via a dichroic mirror followed by a band pass filter of 596.5-667.5 nm wavelength. Pre-processing and further data analysis was performed offline using custom written scripts in MATLAB.

### Pharmacology

Microiontophoresis of MDL-100907 (Sigma-Aldrich; 5-HT2A antagonist, 20 mM in 0.9% NaCl PH 10) or WAY-100135 (Sigma-Aldrich; 5-HT1A antagonist, 5 mM in 0.9% NaCl PH 4) –Although, WAY has been found to be an agonist for 5-HT1B and 5-HT1D receptors (Davidson et al., 1997) as well, it still the best antagonist to use for 5-HT1A receptors– was performed via a pipette inserted to the brain tissue in the region of interest (ROI). ROI was defined beforehand as the region within V1 that showed highest activity levels in response to a short (200 ms) visual stimulus recorded over 3 seconds. A small craniotomy was then made within the ROI and the pipette filled with MDL or WAY solutions was inserted 300 μm below the surface of the brain. Microiontophoresis was performed concurrently with RCaMP imaging and DRN photostimulation keeping all conditions unaltered. Drug solution was retained in the pipette by applying a −10 nA retention current using a constant-current pump (Union-40 microiontophoresis pump; Kation Scientific). To apply the drugs, ejection current was delivered at 60 nA one second before onset of DRN photostimulation, and constantly delivered for 18 seconds. Similar to all other conditions the interval between conditions was 60 seconds. We noticed that in each trial the response to the first visual stimulus of the stimulus train was identical (also to controls), indicating that the inter-trial interval was of sufficient time to allow abolishment of antagonists between each trial. Control experiments were done by microiontophoresis of 0.9% of NaCl solution with a pH of 4.

### In vivo extracellular multi-unit recording

Extracellular multi-unit recordings were performed using tungsten electrodes (0.127 mm diameter, 1 MΩ, WPI, FL, USA), at cortical depths between 250 and 750 μm. Neuronal activity was amplified 1,000× and band-pass filtered between 2 and 5 kHz using an amplifier of Thomas Recording, Germany. The signal was recorded at a sampling rate of 40 kHz using a CED Micro1401 controlled by the Spike 2 software (Cambridge Electronics Design, Cambridge, UK). Spike detection was performed with the Spike 2 software, using a threshold above and/or below the baseline.

Traces of spike counts were averaged over time (200 ms bins) and across trials for each recorded condition. To normalize the data average spike count over 1 second after onset of recordings was subtracted from each individual trace. For further normalization, the traces for each condition were divided by the response amplitude to the first stimulus in the visually-evoked condition. The normalized traces were then averaged over the entire population of cells.

The evoked component of the photostimulated condition (V_ph_) in Figure 4aiv was isolated through subtraction of the spontaneous component (S_ph_) before baseline subtraction. The amplitude of the evoked responses (Figure 4bi) was calculated from the normalized traces around response peak time, i.e., in the time bin 200 ms after the onset of each visual stimulus. Baseline calculations from the normalized traces (Figure 4bii) were performed over time window of 1 second before each of the visual stimulus presentations (Figure 4bii, black, gray, dark blue). The same time windows were used for calculation of spontaneous activity levels during photostimulation (Figure 4bii, light blue).

### Immunohistochemistry

Immunohistochemical localization of 5-HT2A receptors in the visual cortex was performed free-floating using the following primary antibodies: goat anti 5-HT2A receptor antibody (sc15074, dilution 1:100, Santa Cruz Biotechnology), mouse anti parvalbumin antibody (P3038, dilution 1:500, Sigma), mouse anti somatostatin (sc-74556, dilution 1:100, Santa Cruz Biotechnology), rabbit anti GluR2/3 (dilution 1:300, Millipore 07-598). Mouse brains were postfixed for 2 hours after perfusion with 4% PFA in PBS. Coronal sections (30 μM) were collected in 24-well plates in tris-buffered saline (TBS, pH 7.5). Sections were rinsed three times in TBS and subsequently blocked with 0.1% TBST (TBS + Triton X-100) with 3% NDS for one hour at room temperature. The blocking serum was aspirated, and sections were incubated overnight at 4 °C on an orbital shaker with primary antibodies diluted in 1.5% NDS in 0.1% TBST. Brain slices were washed with TBS three times and incubated with anti–species-specific secondary antibodies (DyLight 488 donkey anti rabbit, Thermofisher, Alexa Fluor 568 donkey anti mouse, Alexa Fluor 633 donkey anti goat, Life Technologies, 1:500) in 1.5 % NDS in 0.1 % TBST for one hour at room temperature. Sections were mounted onto Superfrost/Plus Microscope Slides (Thermo Scientific) and coverslipped using Roti-Mount FluorCare (Carl Roth).

Tracer studies (Figure 1b and c) were performed via pressure injection of double-floxed tdTomato virus (AAV2/1.CAG.FLEX.tdTomato.WPRE.bGH; Penn Vector Core) into the DRN of ePet-Cre mice at −4.5 AP to Bregma. Identification of tracer injection and projection fibers in combination with tryptophan-hydroxylase immunohistochemistry (mouse anti TPH antibody, dilution 1:200; Sigma-Aldrich with secondary antibody Alexa Fluor 488 donkey anti mouse) was performed after four weeks of virus expression. Green fluorescent Nissl stain (NeuroTrace 500/525; Life Technologies) was used to provide an anatomic overview of nuclear boundaries.

Immunohistochemical analysis of virus expression in combination with cFos staining after optrode recordings in the DRN were conducted with rabbit anti cFos antibody (sc-52, dilution 1:1000, Santa Cruz Biotechnology) in combination with secondary antibody Alexa Fluor 488 donkey anti rabbit, dilution 1:500, Life Technologies).

Digital images were acquired from brain sections using a Leica TCS SP5 confocal laser scanning microscope interfaced to a personal computer running Leica Application Suite AF 2.6 software. Objectives of 10×/0.3 NA, 20×/0.7 NA were used to capture images. Sequential Z-stacks were created for each section. Captured images were transferred into ImageJ 1.45s (National Institutes of Health) for processing and image overlay.

### Data analysis

Imaging signals were averaged across trials and to remove differences in noise levels due to spatial inhomogeneity in illumination, pixels were divided by pre-stimulus average activity, comprising 200 ms after onset of recording. Furthermore, average blank conditions (see above) were subtracted from all conditions. Therefore, noise due to heartbeat and breathing artefacts were roughly removed after averaging all the trials. Dividing the outcome to the mean blank signal leads to a unit less relative signal of fluorescence, denoted by Δf/f, that make the pre-processed signal independent from global fluctuations occurring during the experiment. In a second step ICA (Maeda et al., 2001; Spors and Grinvald, 2002) was used to remove residual heartbeat and respiratory noise and also for the removal of the photostimulation artefact with a distinct 20 Hz frequency component. These steps were applied to each trial separately and then averaged. The resulting averaged signals were smoothed using a two-dimensional Gaussian filter (σ =20 pixels) and a high pass Butterworth filter (order 4, cut off frequency σ = 33 pixels). The ROI was selected visually in which it’s encircling pixels on the high activities in pixels comprising V1. All spatially averaged traces shown in this paper are averages across pixels in the selected ROI and each response trace was normalized to the first peak in response to the first visual stimulus in control conditions and then averaged across experiments and mice.

Latencies of evoked responses were calculated by finding the time points at which normalized activity crossed 10% of peak response. Response duration was calculated as the time window during which rising and falling edges of the evoked response were above 10% of peak response.

Evoked responses to different visual contrasts were normalized to the peak of the evoked response to 100% grating contrast. Peak maxima of each contrast response (mean over experiments) were fitted using the Naka-Rushton function (Naka and Rushton, 1966):

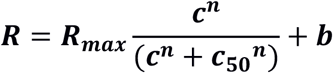

in which *c* is the fractional contrast, c_50_ is the contrast at which its response is half of the maximum response, R_max_ is the maximum response and b is the offset of responses R.

### Statistical analysis

Two-tailed t-test and run length corrected permutation test (Appelbaum et al., 2006; Rekauzke et al., 2016) as a randomization method were used to assess the significance of the results in Figure. 3, Figure 5 and Figure 6. Wilcoxon signed-rank test was used to assess significance of the results shown in Figure 4 and Figure 6 – Figure supplements 6-1.

## Acknowledgments

This work was financially supported by grants from the Deutsche Forschungsgemeinschaft, JA 945/5-1, MA 5806/1-2, MA 5806/2-1, SPP 1665 (JA 945/4-1, HE 2471/12-1), SPP 1926 (HE 2471/18-1) the SFB-874 (A2 & B10), the SFB-1280 (A07) and by the German-Israeli Project Cooperation (DIP, JA 945/3-1, SL 185/1-1). We thank Drs. Knut Holthoff and Knut Kirmse for helpful technical advices on GCaMP imaging.

## Competing interests

The authors declare no competing financial interests.

## References

Adesnik H, Scanziani M (2010) Lateral competition for cortical space by layer-specific horizontal circuits. Nature 464:1155–1160

Akerboom J et al. (2013) Genetically encoded calcium indicators for multi-color neural activity imaging and combination with optogenetics. Front Mol Neurosci 6:2

Allen WE, Kauvar I V, Chen MZ, Richman EB, Yang SJ, Chan K, Gradinaru V, Deverman BE, Luo L, Deisseroth K (2017) Global representations of goal-directed behavior in distinct cell types of mouse neocortex. Neuron 94:891–907.e6

Angelucci A, Bressloff PC (2006) Contribution of feedforward, lateral and feedback connections to the classical receptive field center and extra-classical receptive field surround of primate V1 neurons. Progress in brain research 154: 93–120

Appelbaum LG, Wade AR, Vildavski VY, Pettet MW, Norcia AM (2006) Cue-invariant networks for figure and background processing in human visual cortex. J Neurosci 26:11695–11708

Atallah BV, Bruns W, Carandini M, Scanziani M (2012) Parvalbumin-expressing interneurons linearly transform cortical responses to visual stimuli. Neuron 73:159–170

Carandini M, Heeger DJ (2011) Normalization as a canonical neural computation. Nat Rev Neurosci 13:51–62

Celada P, Puig MV, Artigas F (2013) Serotonin modulation of cortical neurons and networks. Front Integr Neurosci 7:25

Dana H, Mohar B, Sun Y, Narayan S, Gordus A, Hasseman JP, Tsegaye G, Holt GT, Hu A, Walpita D, Patel R, Macklin JJ, Bargmann CI, Ahrens MB, Schreiter ER, Jayaraman V, Looger LL, Svoboda K, Kim DS (2016) Sensitive red protein calcium indicators for imaging neural activity. elife 5:e12727

Davidson C, Ho M, Price GW, Jones BJ, Stamford JA (1997) (+)-WAY 100135, a partial agonist, at native and recombinant 5-HT _1B/1D_ receptors. Br J Pharmacol 121:737–742

Davis M, Strachan DI, Kass E (1980) Excitatory and inhibitory effects of serotonin on sensorimotor reactivity measured with acoustic startle. Science 209:521–523

Dayan P (2012) Twenty-five lessons from computational neuromodulation. Neuron 76:240–256

Dayan P, Abbott LF (2001) Theoretical neuroscience: computational and mathematical modeling of neural systems. Massachusetts Institute of Technology Press.

Doya K (2008) Modulators of decision making. Nat Neurosci 11:410–416

Dugué GP, Lörincz ML, Lottem E, Audero E, Matias S, Correia PA, Léna C, Mainen ZF (2014) optogenetic recruitment of dorsal raphe serotonergic neurons acutely decreases mechanosensory responsivity in behaving mice. PLoS One 9: e105941

Dyck R, Cynader M (1993) Autoradiographic localization of serotonin receptor subtypes in cat visual cortex: transient regional, laminar, and columnar distributions during postnatal development. J Neurosci 10:4316–4338

Franklin KBJ, Paxinos G (2001) Paxinos and Franklin’s The mouse brain in stereotaxic coordinates. Academia press

Gasanov GG, Mamedov ZG, Samedova NF (1989) Changes in reactivity of neurons of the visual cortex under influence of the posterolateral hypothalamus and the nuclei of the midbrain raphe. Neurosci Behav Physiol 19:169–175

Geyer M, Vollenweider F (2008) Serotonin research: contributions to understanding psychoses. Trends Pharmacol Sci 29:445–453

Hale MW, Lowry CA (2011) Functional topography of midbrain and pontine serotonergic systems: implications for synaptic regulation of serotonergic circuits. Psychopharmacology (Berl) 213:243–264

Hannon J, Hoyer D (2008) Molecular biology of 5-HT receptors. Behav Brain Res 195:198–213

Huang L, Yuan T, Tan M, Xi Y, Hu Y, Tao Q, Zhao Z, Zheng J, Han Y, Xu F, Luo M, Sollars PJ, Pu M, Pickard GE, So K-F, Ren C (2017) A retinoraphe projection regulates serotonergic activity and looming-evoked defensive behaviour. Nat Commun 8:14908

Hurley LM, Devilbiss DM, Waterhouse BD (2004) A matter of focus: Monoaminergic modulation of stimulus coding in mammalian sensory networks. Curr Opin Neurobiol 14:488–495

Jakab R, Goldman-Rakic P (1998) 5-Hydroxytryptamine2A serotonin receptors in the primate cerebral cortex: possible site of action of hallucinogenic and antipsychotic drugs in pyramidal cell apical dendrites. Proc Natl Acad Sci USA 95:735–740

Jang H, Cho K, Park S, Kim M, Yoon S (2012) Layer-specific serotonergic facilitation of IPSC in layer 2/3 pyramidal neurons of the visual cortex. J Neurophysiol 107: 407–416

Kim TH, Zhang Y, Lecoq J, Jung JC, Li J, Zeng H, Niell CM, Schnitzer MJ (2016) Long-term optical access to an estimated one million neurons in the live mouse cortex. Cell Rep 17:3385–3394

Kirmse K, Kummer M, Kovalchuk Y,Wite OW, Garaschuk O, Holthof k (2015) GABA depolarizes immature neurons and inhibits network activity in the neonatal neocortex in vivo. Nat Commun 6:7750

Krnjevic K, Phillis JW (1963) Iontophoretic studies of neurones in the mammalian cerebral cortex. J Physiol 165:274–304

Lee S-H, Kwan AC, Zhang S, Phoumthipphavong V, Flannery JG, Masmanidis SC, Taniguchi H, Huang ZJ, Zhang F, Boyden ES, Deisseroth K, Dan Y (2012) Activation of specific interneurons improves V1 feature selectivity and visual perception. Nature 488:379–383

Leysen JE (2004) 5-HT2 receptors. Curr Drug Targets CNS Neurol Disord 3:11–26

Li X, Gutierrez D V, Hanson MG, Han J, Mark MD, Chiel H, Hegemann P, Landmesser LT, Herlitze S (2005) Fast noninvasive activation and inhibition of neural and network activity by vertebrate rhodopsin and green algae channelrhodopsin. Proc Natl Acad Sci U S A 102:17816–17821

Lottem E, Lörincz ML, Mainen ZF (2016) Optogenetic activation of dorsal raphe serotonin neurons rapidly inhibits spontaneous but not odor-evoked activity in olfactory cortex. J Neurosci 36:7–18

Lowery RL, Majewska AK (2010) Intracranial injection of adeno-associated viral vectors. J Vis Exp (45):2140

Lucki I (1998) The spectrum of behaviors influenced by serotonin. Biol Psychiatry 44:151–162

Ma H, Harris S, Rahmani R, Lacefield CO, Zhao M, Daniel AGS, Zhou Z, Bruno RM, Berwick J, Schwartz TH (2014) Wide-field in vivo neocortical calcium dye imaging using a convection-enhanced loading technique combined with simultaneous multiwavelength imaging of voltage-sensitive dyes and hemodynamic signals. Neurophotonics 1:015003

Maeda S, Inagaki S, Kawaguchi H, Song W-J (2001) Separation of signal and noise from in vivo optical recording in Guinea pigs using independent component analysis. Neurosci Lett 302:137–140

Mante V, Bonin V, Carandini M (2008) Functional mechanisms shaping lateral geniculate responses to artificial and natural stimuli. Neuron 58:625–638

Millan M, Marin P, Bockaert J, Mannourylacour C (2008) Signaling at G-protein-coupled serotonin receptors: recent advances and future research directions. Trends Pharmacol Sci 29:454–464

Moyanova S, Dimov S (1986) Modulation of visual excitability cycles in some brain structures by high-frequency stimulation of raphe dorsal nucleus in cats. Acta Physiol Pharmacol Bulg 12:17–25

Muzerelle A, Scotto-Lomassese S, Bernard JF, Soiza-Reilly M, Gaspar P (2016) Conditional anterograde tracing reveals distinct targeting of individual serotonin cell groups (B5–B9) to the forebrain and brainstem. Brain Struct Funct 221:535–561

Naka KI, Rushton WA (1966) S-potentials from luminosity units in the retina of fish (Cyprinidae). J Physiol 185:587–599

Petzold GC, Hagiwara A, Murthy VN (2009) Serotonergic modulation of odor input to the mammalian olfactory bulb. Nat Neurosci 12:784–791

Pinto L, Goard MJ, Estandian D, Xu M, Kwan AC, Lee S-H, Harrison TC, Feng G, Dan Y (2013) Fast modulation of visual perception by basal forebrain cholinergic neurons. Nat Neurosci 16:1857–1863

Pollak Dorocic I, Fürth D, Xuan Y, Johansson Y, Pozzi L, Silberberg G, Carlén M, Meletis K (2014) A whole-brain atlas of inputs to serotonergic neurons of the dorsal and median raphe nuclei. Neuron 83:663–678

Porciatti V, Pizzorusso T, Maffei L (1999) The visual physiology of the wild type mouse determined with pattern VEPs. Vision Res 39:3071–3081

Portas C, Bjorvatn B, Ursin R (2000) Serotonin and the sleep/wake cycle: special emphasis on microdialysis studies. Progress in neurobiology 60:13–35

Ranade SP, Mainen ZF (2009) Transient firing of dorsal raphe neurons encodes diverse and specific sensory, motor, and reward events. J Neurophysiol 102:3026–3037

Reader TA (1978) The effects of dopamine, noradrenaline and serotonin in the visual cortex of the cat. Experientia 34:1586–1588

Rekauzke S, Nortmann N, Staadt R, Hock HS, Schoner G, Jancke D (2016) Temporal asymmetry in dark-bright processing initiates propagating activity across primary visual cortex. J Neurosci 36:1902–1913

Riga M, Bortolozzi A, Campa L, Artigas F, Celada P (2016) The serotonergic hallucinogen 5-methoxy-N, N-dimethyltryptamine disrupts cortical activity in a regionally-selective manner via 5-HT1A and 5-HT2A receptors. Neuropharmacology 101: 370–378

Roerig B, Katz LC (1997) Modulation of intrinsic circuits by serotonin 5-HT3 receptors in developing ferret visual cortex. J Neurosci 17:8324–8338

Santana N, Bortolozzi A, Serrats J, Mengod G, Artigas F (2004) Expression of serotonin1A and serotonin2A receptors in pyramidal and GABAergic neurons of the rat prefrontal cortex. Cereb Cortex 14:1100–1109

Sato TK, Haider B, Häusser M, Carandini M (2016) An excitatory basis for divisive normalization in visual cortex. Nat Neurosci 19:568–570

Sato TK, Häusser M, Carandini M (2014) Distal connectivity causes summation and division across mouse visual cortex. Nat Neurosci 17:30–32

Scott MM, Wylie CJ, Lerch JK, Murphy R, Lobur K, Herlitze S, Jiang W, Conlon RA, Strowbridge BW, Deneris ES (2005) A genetic approach to access serotonin neurons for in vivo and in vitro studies. Proc Natl Acad Sci U S A 102:16472–16477

Seillier L, Lorenz C, Kawaguchi K, Ott T, Nieder A, Pourriahi P, Nienborg H (2017) Serotonin decreases the gain of visual responses in awake Macaque V1. J Neurosci 37:11390–11405

Seybold BA, Phillips EAK, Schreiner CE, Hasenstaub AR (2015) Inhibitory actions unified by network integration. Neuron 87:1181–1192

Shimegi S, Kimura A, Sato A, Aoyama C, Mizuyama R, Tsunoda K, Ueda F, Araki S, Goya R, Sato H (2016) Cholinergic and serotonergic modulation of visual information processing in monkey V1. J Physiol 110:44–51

Shukla R, Watakabe A, Yamamori T (2014) mRNA expression profile of serotonin receptor subtypes and distribution of serotonergic terminations in marmoset brain. Frontiers in neural circuits 8: 52

Soubrié P (1986) Reconciling the role of central serotonin neurons in human and animal behavior. Behav Brain Sci 9:319

Spors H, Grinvald A (2002) Spatio-temporal dynamics of odor representations in the mammalian olfactory bulb. Neuron 34:301–315

Tian L, Hires SA, Mao T, Huber D, Chiappe ME, Chalasani SH, Petreanue L, Akerboom J, Mckinney SA, Schreiter ER, Bargmann CI, Jayaraman V, Svoboda K, Looger LL (2009) Imaging neural activity in worms, flies and mice with improved GCaMP calcium indicators. Nat Methods 6:875–881

Tyree S, de Lecea L (2017) Optogenetic investigation of arousal circuits. International Journal of Molecular Sciences 18:1773.

Urban DJ, Zhu H, Marcinkiewcz CA, Michaelides M, Oshibuchi H, Rhea D, Aryal DK, Farrell MS, Lowery-Gionta E, Olsen RHJ, Wetsel WC, Kash TL, Hurd YL, Tecott LH, Roth BL (2016) Elucidation of the behavioral program and neuronal network encoded by dorsal raphe serotonergic neurons. Neuropsychopharmacology 41:1404–1415

Wallace DJ, zum Alten Borgloh SM, Astori S, Yang Y, Bausen M, Kügler S, Palmer AE, Tsien RY, Sprengel R, Kerr JND, Denk W, Hassan MT (2008) Single-spike detection in vitro and in vivo with a genetic Ca2+ sensor. Nat Metohds 5:797–804.

Watakabe A, Komatsu Y, Sadakane O, Shimegi S, Takahata T, Higo N, Tochitani S, Hashikawa T, Naito T, Osaki H, Sakamoto H, Okamoto M, Ishikawa A, Hara S, Akasaki T, Sato H, Yamamori T (2009) Enriched expression of serotonin 1B and 2A receptor genes in macaque visual cortex and their bidirectional modulatory effects on neuronal responses. Cereb Cortex 19:1915–1928

Waterhouse BD, Ausim Azizi S, Burne RA, Woodward DJ (1990) Modulation of rat cortical area 17 neuronal responses to moving visual stimuli during norepinephrine and serotonin microiontophoresis. Brain Res 514:276–292

Weber ET, Andrade R (2010) Htr2a gene and 5-HT(2A) receptor expression in the cerebral cortex studied using genetically modified mice. Front Neurosci 4:36

Wilson NR, Runyan CA, Wang FL, Sur M (2012) Division and subtraction by distinct cortical inhibitory networks in vivo. Nature 488:343–348

Xiao D, Vanni MP, Mitelut CC, Chan AW, LeDue JM, Xie Y, Chen AC, Swindale N V, Murphy TH (2017) Mapping cortical mesoscopic networks of single spiking cortical or sub-cortical neurons. elife 6:e19976

Zhang G, Stackman RW (2015) The role of serotonin 5-HT2A receptors in memory and cognition. Front Pharmacol 6:225

